# Seasonal tissue-specific gene expression reveals reproductive and stress-related transcriptional systems in wild crown-of-thorns starfish

**DOI:** 10.1101/2023.12.18.572276

**Authors:** Marie Morin, Mathias Jönsson, Conan K. Wang, David J. Craik, Sandie M. Degnan, Bernard M. Degnan

## Abstract

Animals are influenced by the season, yet we know little about the changes that occur in most species throughout the year. This is particularly true in tropical marine animals that experience relatively small annual temperature and daylight changes. Like many coral reef inhabitants, the crown-of-thorns starfish (COTS), well known as a notorious consumer of corals and destroyer of coral reefs, reproduces exclusively in the summer. By comparing gene expression in seven somatic tissues procured from wild COTS sampled on the Great Barrier Reef, we identified more than 2,000 protein-coding genes that change significantly between summer and winter. COTS genes that appear to mediate conspecific communication – including both signalling factors released into the surrounding sea water and cell surface receptors – are upregulated in external secretory and sensory tissues in the summer, often in a sex-specific manner. The sexually dimorphic gene expression appears to be underpinned by sex- and season-specific transcription factors (TFs) and gene regulatory programs. There are over 100 TFs that are seasonally expressed, 86% of which are significantly upregulated in the summer. Six nuclear receptors are upregulated in all tissues in the summer, suggesting that systemic seasonal changes are hormonally controlled, as occurs in vertebrates. Unexpectedly, there is a suite of stress-related chaperone proteins and TFs, including HIFα, ATF3, C/EBP, CREB and NF-κB, that are uniquely co-expressed in gravid females. The upregulation of these stress proteins in the summer suggests the demands of oogenesis in this highly fecund starfish affects protein stability and turnover in somatic cells. Together, these circannual changes in COTS gene expression provide novel insights into seasonal changes in this coral reef pest and have the potential to identify vulnerabilities for targeted biocontrol.

## Introduction

The crown-of-thorns starfish (COTS, *Acanthaster planci* species complex) is a predator of reef-building corals that lives in the oceans of the Indo-Pacific [1–5]. The very high seasonal fecundity of this starfish, along with the timing of its spawning and the high dispersal and survival potential of its larvae, appear to contribute to population fluctuations and outbreaks that cause extensive loss of coral cover and associated reef biodiversity [3,4,6–16].

COTS, as is the case with many echinoderms, reproduce by broadcast spawning in aggregations that form in the summer in response to conspecific and environmental cues [8,10,11,17–20]. Understanding the molecular basis of the conspecific communication required for aggregation and spawning could provide novel ways to disrupt the starfish reproduction and thus mitigate destructive outbreaks [21]. As gene expression levels can provide insights into the regulatory and physiological state of an organism [22–26], transcriptomic analyses have the potential to identify genes underlying the mechanisms used by COTS to adjust their physiology and behaviours according to time of year. For instance, gravid males and females differentially express genes encoding signalling factors released into the surrounding sea water and chemosensory receptors that potentially detect these signals [21,28].

To characterise how COTS change seasonally, we assessed gene expression in seven somatic tissues isolated immediately after the starfish were hand-collected on the Great Barrier Reef in the summer and winter. This sampling strategy allowed us to identify genes that are differentially expressed between gravid, aggregating female and male starfish prior to summer spawning.

## Results

### Sampling wild crown-of-thorns starfish on the Great Barrier Reef

We analysed CEL-Seq2 transcriptomes generated from RNA procured from concentrated coelomocytes, and small biopsies of papulae, radial nerve, sensory tentacles, skin, spines and tube feet from 20 individual COTS removed by hand from Davies and Lynch’s Reefs on the Great Barrier Reef (Fig 1A and S1 Table; see Materials and Methods and [28] for details). Seven females and six males were sampled in the summer, and seven unsexed individuals in the winter (gonads were not visible). All COTS collected in the summer were gravid, indicating that a spawning event was imminent [8,17,28,]. All individuals were dissected within two hours of removal from the reef and maintained in ambient seawater on a vessel anchored at the collection site until dissection. At the times of field collection, the mean summer and winter sea surface temperatures were 27.7 and 23.0°C, respectively, which are within the normal seasonal range (Fig 1B). Day lengths, as measured from sunrise to sunset, were approximately 13.2 and 11.3 hours in the summer and winter, respectively. There were no notable weather events prior to, or at the time of, sampling.

**Fig 1.**
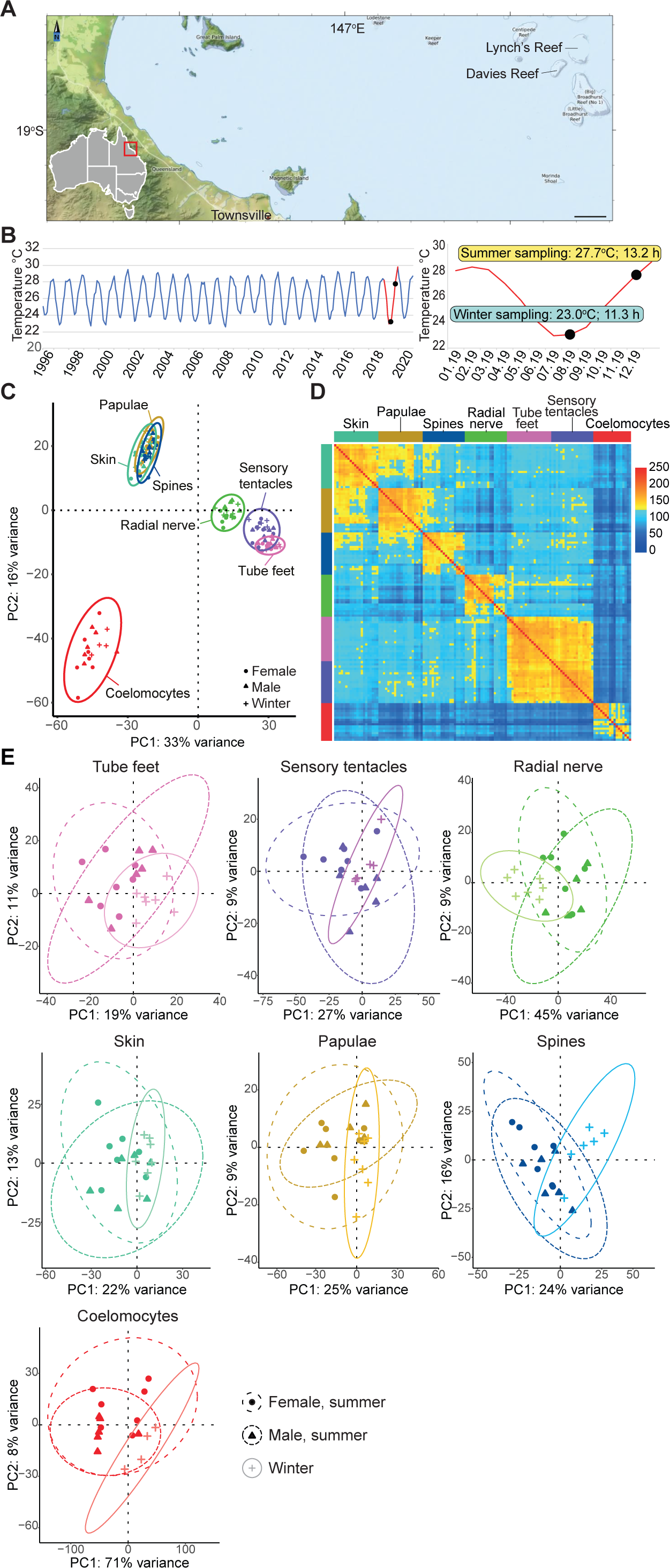
Gene expression in wild COTS. **(A)** Field sites where RNA was procured from COTS tissues in 2019. **(B)** Sea temperatures at 4 m depth at Davies Reef from 1996 to 2020. 2019 is highlighted in red and the times of COTS sampling are marked with black dots. Water temperature and day length (sunrise to sunset) at sampling times are shown. Map and temperature data were obtained from the Australian Institute of Marine Science. **(C)** PCA of tissue transcriptomes; 95% confidence ellipses shown. **(D)** Hierarchical clustered heatmap of Pearson correlation of expressed genes across the seven tissue transcriptomes. **(E)** PCAs with 95% confidence ellipses, showing the differences in overall gene expression between males and females (summer), and unsexed COTS (winter). Pink, tube feet; purple, sensory tentacles; green, radial nerve; turquoise, skin; brown, papulae; blue, spines; red, coelomocytes; circles, females; triangles, males; crosses, unsexed winter individuals.

Although we procured small biopsies from specific regions of the external tissues (e.g. single tube foot per individual) and restricted the coelomic fluid to a few ml (see Materials and Methods), we acknowledge that the transcriptomes nonetheless are likely to be derived from multiple cell types. However, for simplicity we refer to these as “tissues” throughout this report.

### Somatic tissue gene expression

On average, 69.7% of the CEL-Seq2 reads from the 134 tissue transcriptomes that passed quality control filtering mapped to the GBR v1.1 gene models (see Materials and Methods) [29], with each tissue expressing on average 15,490 protein-coding mRNAs (Table 1 and S1, S2 Tables). Principal component analysis (PCA) and hierarchical clustering analysis revealed that gene expression in each tissue is very similar between individuals, regardless of sex or season (Fig 1C, D). This affirmed that the sampling regime employed yielded consistent tissue-specific gene expression profiles.

**Table 1.** Number of expressed protein coding sequences in COTS tissues.

|  | **Total**<br>**expressed** | **Significantly**<br>**upregulated<sup>1</sup>** | **Uniquely**<br>**expressed<sup>2</sup>** |
| --- | --- | --- | --- |
| Tube feet | 15,771 | 266 | 79 |
| Sensory tentacles | 16,256 | 222 | 185 |
| Radial nerve | 14,226 | 772 | 23 |
| Skin | 15,842 | 372 | 115 |
| Papulae | 16,416 | 1,356 | 307 |
| Spines | 15,779 | 288 | 153 |
| Coelomocytes | 14,137 | 1,386 | 79 |

<sup>1</sup> Protein coding mRNAs significantly upregulated compared to all other tissues (DESeq2, adjusted *p*-value < 0.05)
<sup>2</sup> Protein coding mRNAs not detected in any other tissue

The transcriptomes of oral (downwardly-facing) radial nerve, sensory tentacles and tube feet are most similar to each other and are distinct from those of the aboral (upwardly-facing) papulae, skin and spines, which are also most similar to each other (Fig 1C, D). The coelomocytes, the only internal tissue analysed, is distinct from both oral and aboral tissues. Pairwise comparisons of gene expression using DESeq2 identified between 222 and 1,386 protein-coding mRNAs that are significantly differentially expressed within a given tissue (adjusted *p*-value < 0.05; Table 1 and S3 Table). The three oral tissues together uniquely upregulate 1,752 coding sequences that are enriched in neural and sensory functions, consistent with the known biological roles of these tissues [30–34]; 71.9% of these genes are upregulated in only one tissue (Table 1, S3 and S4 Table). In contrast, 2,786 coding sequences upregulated in aboral tissues are enriched in immune and cilia function, again consistent with known roles of these tissues [35,36]; 72.3% of these are uniquely upregulated in only one tissue (Table 1, S3 and S4 Table). Nearly 1,400 coding sequences are upregulated in the internal coelomocytes compared to the six external tissues (Table 1 and S3 Table).

### Seasonal changes in gene expression

Like most marine animals, COTS are poikilotherms (ectotherms) and thus their metabolism and physiology is strongly influenced by the surrounding water temperature [37,38]. COTS inhabiting Davies and Lynch’s Reefs in the central part of the Great Barrier Reef typically experience 6-7°C annual change in water temperature and a maximum of 2.3 hours difference in day length (sunrise to sunset on summer and winter solstice). These environmental variations are sufficient for COTS and a diversity of coral reef animals to seasonally change their reproductive status, with COTS typically spawning in the austral summer from November to January [8,10,17,19]. The exact timing of spawning appears to be based on a combination of abiotic and biotic cues [10,17,18,20,21,28]. At the time we undertook the summer sampling (December 2019), all COTS were gravid and appeared to have not yet spawned, consistent with previous observations of the reproductive status of this starfish on Davies Reef [17]. In contrast, gonads were not visible in any individuals sampled in the winter (August 2019) and thus their sex or possible level of hermaphrodism [39] could not be determined. In any case, winter transcriptomes clustered by tissue type with male and female summer transcriptomes, indicating that tissue type was the greatest determinant of transcriptional state in both seasons, regardless of sex (Fig 1C, E).

We identified 2,079 protein-coding mRNAs significantly differentially expressed between seasons in at least one tissue (adjusted *p*-value < 0.05); 71.8% of these were upregulated in summer (Table 2). Protein processing pathways were more activated in most tissues in summer, consistent with the upregulation of multiple protein chaperones, cell surface receptors (including 29 G-protein coupled receptors, GPCRs) and secreted proteins that include conserved hormones, neuropeptides and developmental proteins (Fig 2A, B and S5 Table). Notably, the external sensory tissues – radial nerve, spines and tube feet – had the greatest seasonal differences in gene expression (Fig 2C). In the radial nerve, this includes the upregulation of components of Wnt, FoxO, AGE-RAGE and phosphatidylinositol signalling pathways in the summer, consistent with a change in physiological and cell states, and an increase in overall intercellular signalling (Fig. 2A, B).

**Fig 2.**
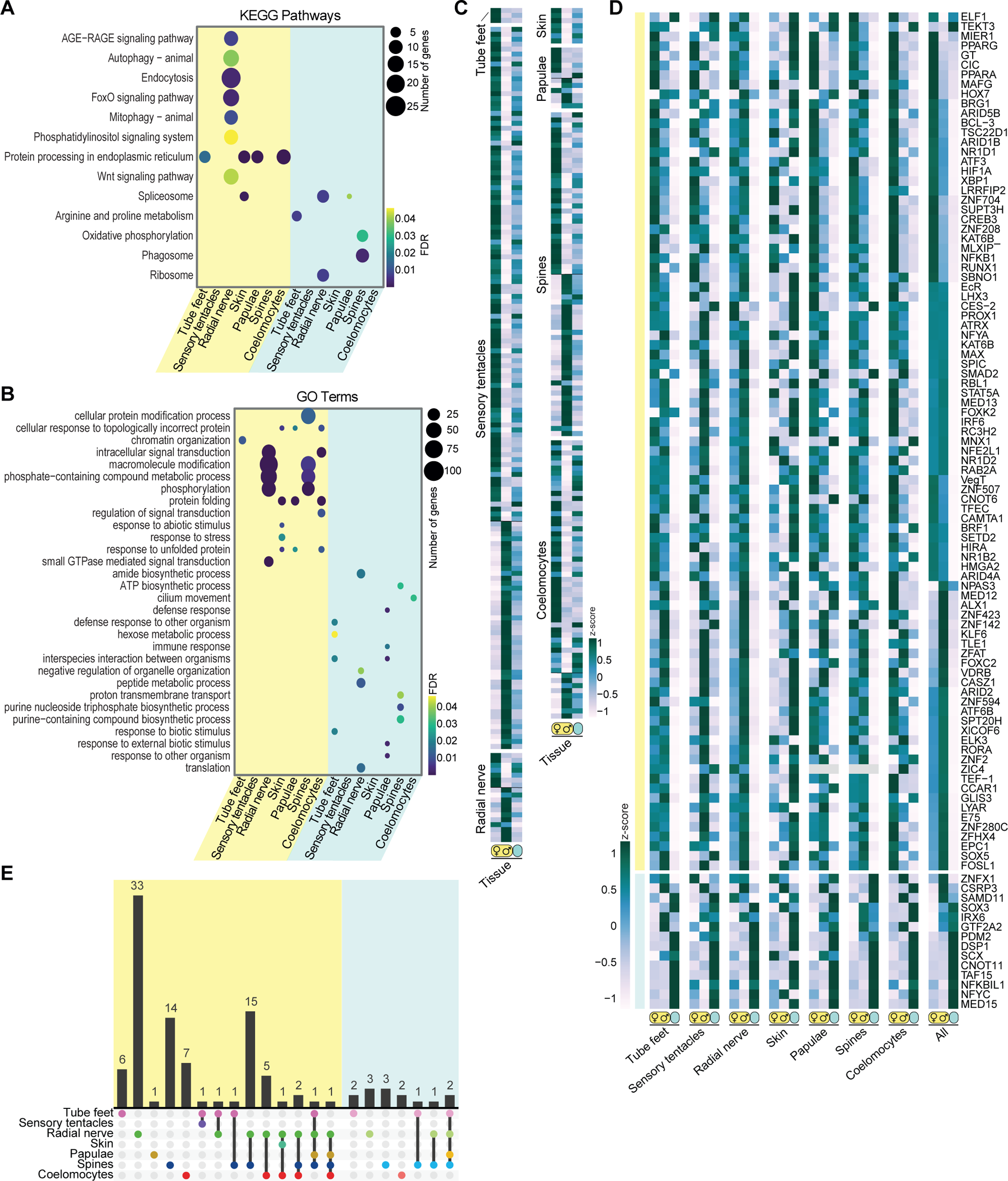
Differential gene expression between seasons and sexes. **(A)** KEGG pathways enriched in summer (yellow) and winter (blue) upregulated coding sequences in each tissue. **(B)** GO enrichments for upregulated coding sequences in the summer and winter for each tissue. In **(A)** and **(B)**, the dot size and colour corresponds to the number of coding sequences associated with each term and the false discovery rate (FDR)-corrected *p*-value, respectively. **(C)** Heatmap of differentially expressed coding sequences in gravid female and male COTS prior to spawning. Scaled (z-score) expression levels based on TPM normalised reads from seven females (♀) and six males (♂) in the summer (yellow oval), and seven unsexed individuals in the winter (blue oval). The heatmap is grouped into the tissues where these genes are differentially expressed. **(D)** Heatmap of differentially expressed TFs in wild COTS for each tissue. The “All” heatmap shows the average expression across all tissues across both sexes and seasons. TFs upregulated in the summer (yellow) and winter (blue) are at the top and bottom of the heatmap, respectively. **(E)** Overlap of summer-upregulated (left, yellow) and winter-upregulated (right, blue) TFs in each tissue. The histogram represents the number of differentially expressed TFs shared by one or more tissues, as indicated by the bottom legend.

**Table 2.** Number of significantly upregulated coding sequences in COTS tissues between summer and winter.

|  | Summer |  |  | Winter<sup>2</sup> |  |  |
| --- | --- | --- | --- | --- | --- | --- |
|  | Female | Male | Both<sup>1</sup> | Female | Male | Both<sup>3</sup> |
| Tube feet | 507 | 369 | 169 | 344 | 327 | 126 |
| Sensory tentacles | 151 | 101 | 21 | 53 | 66 | 3 |
| Radial nerve | 1,427 | 1,735 | 980 | 534 | 551 | 200 |
| Skin | 222 | 64 | 35 | 162 | 100 | 7 |
| Papulae | 374 | 79 | 52 | 193 | 40 | 14 |
| Spines | 1,649 | 956 | 664 | 927 | 562 | 302 |
| Coelomocytes | 304 | 632 | 189 | 129 | 910 | 82 |

<sup>1</sup> Upregulated in female and male COTS in the summer
<sup>2</sup> Upregulated in unsexed COTS in the winter relative to either female or male COTS in the summer
<sup>3</sup> Upregulated in unsexed COTS in the winter relative to both female and male COTS in the summer

We previously found that exoproteomes released into the sea water by aggregating COTS are enriched in lineage-specific sequences, such as COTS-specific ependymins (S1 Fig and S6 Table) [21]. In the present study, we found 285 lineage-specific protein coding genes that are differentially expressed between summer and winter. Of these, 202 are upregulated in the summer, including 16 genes that encode proteins secreted by aggregating COTS and seven genes that are expressed in a sex-specific manner (S1 Fig and S6 Table) [21,28]. Notably, we found that spines differentially express the largest number of lineage-specific genes (155), consistent with previous observations that spines and skin both secrete many proteins found in surrounding sea water exoproteomes [21].

### Sex-specific gene regulation in the summer

We recently identified 183 and 100 protein-coding mRNAs that are differentially upregulated respectively in female and male somatic tissues just prior to summer spawning [28]. Here, we found evidence that these genes are seasonally regulated in a sex-specific manner, with relative winter transcript levels being between those of the two sexes in the summer (Fig 2C and S5 Table). These gene expression profiles are consistent with systemic sex-specific changes in physiological state as environmental cues, including increasing water temperature and day length, trigger a preparation for reproduction. Most of these genes, which appear to be related to the onset of reproduction, are expressed in the sensory tentacles and spines, both of which are implicated in conspecific communication during summer aggregation and in the regulation of spawning [28]. We also detected 21 class A rhodopsin-like GPCRs (putative chemoreceptors), 189 secreted proteins and 285 lineage-specific genes that are differentially upregulated in a sex-specific manner in summer (S1 Fig and S6 Table). Together, these findings reveal a starfish-wide upregulation of sex-specific gene expression in the summer that appears to underlie conspecific communication and reproduction.

### Activation of transcription factors in the summer

As part of the general upregulation of genes in the summer, we found that 89 of 102 seasonally-expressed transcription factors (TFs) are significantly upregulated in summer (adjusted *p*-value < 0.05; Fig 2D, E; S5 Table). Although most TFs have unique expression profiles across tissues, sexes and seasons (61 are uniquely upregulated in one specific tissue), the overall expression profile aligns with sex and season. That is, TF expression is higher in either males or females in summer, or is higher in one season compared to the other (Fig 2D). We did not find any significantly upregulated TFs that are shared among all oral tissues or all aboral tissues (Fig 2D), despite the overall transcriptomic similarities of the tissues comprising these two tissue sets (Fig 1C, D). These findings suggest that multiple cell type-specific gene regulatory networks are operating across the somatic cells comprising these tissues. Interestingly, seasonal expression of most TFs in the skin is opposite to other tissues by being relatively higher in the winter (Fig 2D).

Of the 61 TFs that are significantly upregulated in one specific tissue in the summer, 33 and 14 are uniquely upregulated in radial nerves and spines, respectively (Fig 2D). These two functionally different and oppositely facing tissues also have the most TFs in common (15), including three nuclear receptors: nuclear receptor subfamily 1 group D member 1 (NR1D1/Rev-Erbα); nuclear receptor subfamily 1 group D member 2 (NR1D2/ Rev-Erbβ); and peroxisome proliferator-activated receptor gamma (PPARγ) (Fig 2E and S5 Table). The scale and complexity of TF gene expression in spines and the radial nerve suggests that multiple seasonal gene regulatory networks are operating simultaneously across the various cell types that comprise these two somatic tissues.

### Co-expressed summer genes suggest an overall increase in somatic cellular activity and turnover in gravid crown-of-thorns starfish

A weighted gene correlation network analysis (WGCNA) on 9,159 protein coding sequences that met expression and variance thresholds (see Materials and Methods) [40] identified 15 gene co-expression modules that are largely correlated with a specific tissue (S2 Fig and S7 Table). However, three of the co-expression modules – turquoise, cyan and pink – comprise genes that were highly expressed in all tissues in the summer (*p*-values 1e^−3^, 4e^−2^, and 4e^−6^ respectively; Fig 3A, S2 Fig and S7 Table). There were no co-expression modules that comprise genes upregulated in all tissues in the winter.

**Fig 3.**
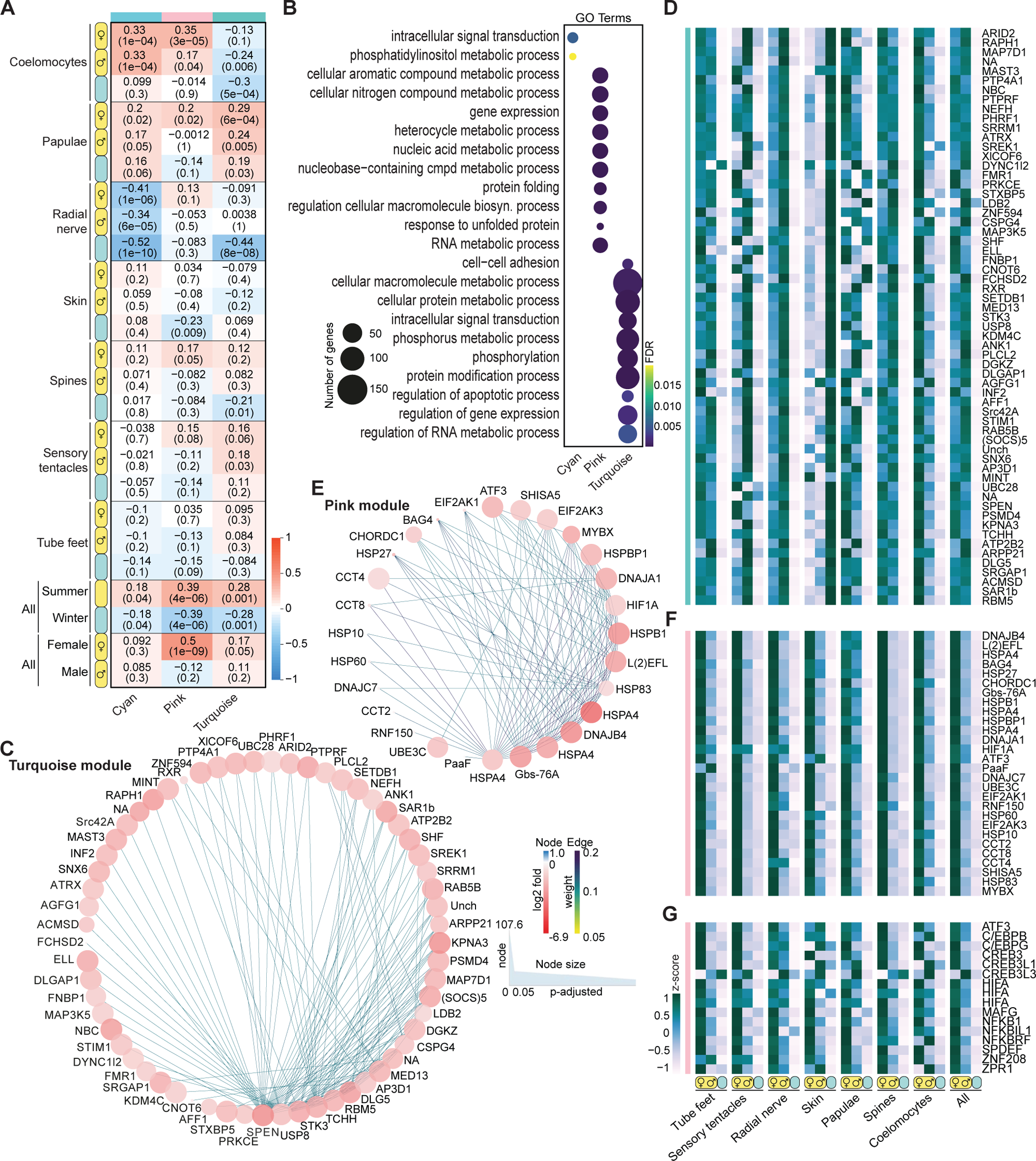
Seasonal gene co-expression. **(A)** Correlation coefficient and *p*-value (in brackets) of co-expressed genes in three modules (cyan, pink, and turquoise) that are comprised of genes that are upregulated in the summer in relation to tissue, season and sex. Red and blue colours depict the strength of the correlation. **(B)** Most significantly enriched biological processes GO terms of the coding sequences comprising cyan, pink and turquoise modules. The dot size reflects the number of coding sequences in each GO term and the colour depicts the FDR-corrected *p*-value. **(C)** Gene interaction networks for the turquoise module (edge weights > 0.164). The size of each coding sequence (node) in the network corresponds to the significance of differential expression (upregulated in the summer; *p*-adjusted value in DESeq2). The colour corresponds to the log2fold change in gene expression: red, upregulated in summer; blue, upregulated in winter. **(D)** Expression profiles of the core hub genes in the turquoise module. **(E)** Gene interaction networks for the pink module (edge weights > 0.164). The size and colour of each coding sequence in the network is as described for the turquoise module. **(D)** Expression profiles of the core hub genes in the pink module. **(G)** Expression profiles of transcription factors identified in the pink module. The heatmaps **(D, F, G)** show the scaled expression levels (z-score) based on TPM normalized reads for each tissue in females, males, and winter individuals. The “All” heatmap shows the average expression across all groups and tissues.

The most interconnected genes in the turquoise and cyan modules have similar expression in both male and female COTS and are enriched in coding sequences involved in gene regulation, signal transduction and enzyme function (Fig 3B-D and S7 Table). This expression profile is consistent with an overall increase in cellular activity and cell turnover in summer compared to winter. The most interconnected gene in this network is a member of the SPEN family of transcriptional repressors, which can interact with nuclear receptors and other corepressors, and contain RNA recognition motifs that can interact with nuclear receptor mRNAs [41,42]. The turquoise module includes six nuclear receptors – retinoid X receptor (RXR), vitamin D3 receptor B (VDR), ecdysone receptor (EcR/N1NH2), NR1D1/Rev-Erbα, NR1D2/ Rev-Erbβ and PPARγ – that may be SPEN targets in somatic tissues of gravid COTS (Fig. 3C, D, S5, and S7 Tables). These nuclear receptors generally are activated by diverse steroid ligands [43], suggesting that summer gene expression profiles are underpinned by a seasonal change in composition of the endogenous ligands (e.g. hormones) to these, and perhaps other constitutively expressed nuclear receptors.

Notably, like the general seasonal TF gene expression profiles in COTS (Fig 2D), all the most interconnected co-expressed upregulated genes in the summer are downregulated in the skin (Fig 3D). This finding suggests that this protective tissue has a markedly different regulatory architecture to the other tissues. Skin shares only one differentially expressed TF gene with the radial nerve and coelomocytes (*ATF3*) and has no uniquely expressed TFs that are significantly differentially expressed between seasons (Fig 2D, E and S5 Table). These gene expression and co-expression profiles provide further evidence that skin changes the least in function of all the tissues between winter and summer, consistent with a year-round role in immunity and defense.

### Female crown-of-thorns starfish upregulate stress proteins in the summer

The pink module (see above) comprises genes that are upregulated in gravid female tissues in the summer *(p-*value 1e^−9^; Fig 3A, S2 Fig and S7 Table). This sex- and season-specific co-expression module is enriched in coding sequences related to protein turnover and stability, and other stress related functions. It includes heat shock proteins (HSPs), hypoxia-inducible factors, and a suite of conserved TFs that are involved in stress responses across the animal kingdom, including bZIP TFs ATF3, C/EBP (CHOP), CREB and MAF, HIFα and NF-κB (Fig 3E-G) [44,45]. The most interconnected genes in this network are conserved heat shock and chaperone proteins, including HSP70 (HSPA), HSP40 (DNAJ), HSP83, HSP60, HSP20 (Protein lethal(2)essential for life), HSP52/27/28 (HSPB1) and HSP10; chaperonin containing TCP1 subunits 2, 4 and 8 (CCT2, 4 and 8); and BAG family molecular chaperone regulator 4. Notably, there is no evidence that the genes in this module, many of which are transiently expressed in response to acute stresses in other animals [45–48], were activated because of a marked change in environmental conditions, such as a marine heat wave or hypoxia on the source reefs (Fig 1A, B). Gravid males collected from the same reef locations and at the same time did not show upregulation of these stress genes. Also significant is that these stress-related genes are upregulated in female skin in the summer, in contrast to other genes in this tissue that are constitutively expressed or downregulated in the summer. Together these findings lead us to conclude that the activation of stress proteins in gravid females reflects an organism-wide state that may relate to the physiological taxing nature of being in a reproductive state.

## Discussion

By comparing gene expression in seven somatic tissues from wild COTS in the summer and winter, we have revealed new insights into seasonal changes in this coral reef pest. The procurement of RNA from small, targeted biopsies from multiple starfish immediately after their removal from the reef yielded consistent gene expression profiles within a given tissue, season and sex, supporting the proposition that this replicated transcriptomic dataset accurately reflects natural gene activity [29]. This is further evidenced by each tissue transcriptome being enriched in protein coding sequences whose roles reflect the known biological function of that tissue. Thus, in addition to identifying differences between seasons and sexes, this approach provides new insights into the natural functioning of tissues and organs in the wild. For instance, compared to other COTS tissues, the radial nerve, sensory tentacles and tube feet upregulate neural and sensory genes, consistent with sensory roles of these organs in COTS and other echinoderms [28, 30–34,36]. This includes the expression of a diversity of chemosensory GCPRs in the most distal sensory tentacles and signalling neuropeptides in the radial nerve in the summer. By contrast, aboral papulae, skin and spines upregulate genes involved in immunity and cilia function, consistent with skin having a general immune function and papulae being comprised of ciliated cells that create countercurrents involved in respiration [35,36]. Unexpectedly, papulae additionally express diverse GPCRs, secreted proteins, TFs and lineage-specific coding sequences, suggesting this tissue has a wider sensory and regulatory role. Coelomocytes isolated from wild COTS are probably derived from the perivisceral coelom, which has a diversity of functions in COTS and other echinoderms, including immunity and wound healing, nutrient transport and waste removal [36,49–51]. We found that the coelomocyte transcriptomes are enriched in genes associated with microfilament function, consistent with phagocytes being a dominant cell type in this tissue as observed in other echinoderms [36,51].

All tissues express a unique repertoire of TFs, with neither oral nor aboral tissues uniquely sharing any TFs to the exclusion of all other tissues, despite their respective transcriptomes being very similar. Surprisingly, the tissues that have the highest diversity of differentially expressed TFs – the radial nerve and spines – also share the most TFs with each other. As these two tissues are functionally different [36], and express transcriptomes of similar complexity to the other tissues, it is unclear why they differentially express a greater diversity of TFs. The presence of a diverse TF repertoire in spines is consistent with this tissue playing a diversity of roles in conspecific communication that include both releasing and receiving external signals [21].

Most of the 2,079 seasonally expressed protein coding genes were significantly upregulated in the summer (71.8%). A subset of these may play a role in conspecific communication underlying pre-spawning aggregations and synchronized spawning events [28], including 21 putative chemosensory GPCRs, 189 secreted proteins, and 285 lineage-specific genes, many of which are expressed in a sex-specific manner. This finding, combined with the differential activation and repression of other sex-specific genes in the summer, suggests that gene activity in the summer is maximally different between sexes, and is likely to underlie behaviours that ensure the pre-spawning formation of male and female aggregations in the wild.

Correlated with the upregulation of conspecific communication and other protein coding genes in the summer are 89 TFs; only 13 TFs are upregulated in the winter. Although most summer TFs are expressed in specific somatic tissues and are thus likely to be part of specific cell and tissue type gene regulatory networks, a subset is co-expressed across all tissues in the summer. These widely expressed TFs may be coordinating the complex tissue-specific transcriptional changes that occur as seawater temperature and day length increases, and as COTS reproductively matures. Amongst these TFs are six nuclear receptors that are known in vertebrates to be influenced by seasonal hormones [43,52]. Putative nuclear receptor ligands, such as steroids, have been detected in COTS and in other echinoderms, and may be playing a seasonal role in reproduction [51,53–55]. These findings are consistent with the existence of conserved deuterostome – or maybe even bilaterian or metazoan – mechanisms to regulate circannual physiological changes via the activation of nuclear receptors and partner TFs.

Unexpectedly, we observed many highly conserved, stress-related chaperones and TFs that are upregulated in somatic tissues of gravid (summer) females, but not males. A single female COTS can produce more than 100 million eggs in one spawning season and the ovary can comprise up to 34% of adult body mass, reflecting the very large energy investment that must be deployed to produce oocytes during the summer reproductive season [7]. On the reef, female COTS are most fecund when sea temperature is near maximal and oxygen levels are lower and have greatest diel fluctuations [56]. As there was no evidence of abnormally high temperatures or other environmental stressors at the time the COTS were sampled, we infer that the upregulation of stress proteins may reflect the overall physiological state of female COTS that are channeling a large proportion of their nutrients to oocytes. It would be interesting to see if this phenomenon occurs in other highly fecund echinoderms and marine animals, as it is currently unclear if this is a general phenomenon used in marine animals that seasonally deploy a large proportion of their resources to the production of gametes.

In conclusion, our atlas of somatic tissue transcriptomes from wild COTS in the summer and winter enables the identification of molecular factors that underpin season- and sex-specific biological processes. Given the large-scale transcriptional changes that occur when COTS are translocated from the wild [29], it appears unlikely that the same depth of insight would be achieved by sampling captive starfish. The seasonal changes in gene expression have revealed a complex, and often sex-specific, interplay between regulatory, signalling and structural genes. Notably, this includes tissue-specific expression profiles that overlap substantially with adjacent or related tissues, and the large-scale activation of genes in the summer, many of which are differentially upregulated in one sex and are involved in cell signalling and conspecific communication. This analysis also uncovered a cryptic physiological state in gravid females that includes a systemic upregulation of conserved stress response genes that may have a protective role prior to spawning. Beyond its utility in understanding the biology of this coral reef pest and providing molecular leads for its future biocontrol, this atlas provides an approach to understand circannual rhythms in tropical and coral reef animals where seasonal differences are less pronounced than elsewhere on Earth.

## Materials and Methods

### Sampling of wild-caught COTS and CEL-Seq2 library construction

Adult *Acanthaster planci* cf. *solaris* (crown-of-thorns starfish; COTS) [1,2] were collected from Davies Reef (18°50’S, 147°39’E) and Lynch’s Reef (18°76’S, 147°63’E) on the Great Barrier Reef (Fig 1A), and tissue RNAs were isolated as previously described [28]. Care was taken to ensure no overlap between tissues such that skin had no papulae and vice versa. Briefly, COTS were collected by hand from the reef and housed in ambient seawater onboard the vessel. Tissues were biopsied and transferred into RNALater less than two hours after the starfish were removed from the reef. Seven unsexed individuals were collected during the winter (August 2019), and seven and six gravid females and males, respectively, were collected during the summer reproductive season (December 2019). Seven somatic tissues – coelomocytes, sensory tentacles, tube feet, radial nerves, skin, papulae and spines – were isolated from each female, male and winter individual, except coelomocytes and spines were not collected from two and one individuals in the winter, respectively.

RNA isolation and quality-assessment, and CEL-Seq2 library construction all were performed as previously described [28]. Tissue from each individual were kept separate throughout the procedures, and each individual sample was uniquely barcoded prior to pooling and sequencing [57]. The CEL-Seq2 libraries were sequenced on an Illumina HiSeq X ten platform at NovogeneAIT Genomics in Singapore.

Raw reads were assessed for quality and adaptor contents using FastQC [58] and analysed using the CEL-Seq2 pipeline (https://github.com/yanailab/CEL-Seq-pipeline; version 1.0) [57]. Reads were trimmed to 35 bp, demultiplexed and mapped to the GBR v1.1 genome using Bowtie2 [29]. Transcript counts of each coding sequences were generated using HTSeq [59]. Samples with mapped transcripts < 0.5 million were discarded as low-quality [60]. GBR v1.1 gene models and transcriptomes can be visualised at https://apolloportal.genome.edu.au/degnan/cots [28,29].

### Identification of lineage-specific classes of expressed genes

Transcription factors and chaperones were identified based on their BLAST2GO annotation [29]. GPCRs were identified by gene classification established by Hall et al. [21]. SignalP 5.0 was used to determine if COTS GBR v1.1 gene models [29] possessed signal peptides and potentially encoded for secreted proteins [61]. These gene models were then assessed for transmembrane domains that would indicate they encoded membrane-bound rather than secreted proteins using TMHMM Server v. 2.0 [62,63]. Proteins were considered secreted if they possessed a predicted signal peptide but no transmembrane domain. Identified secreted proteins were then classified into six categories: hydrolytic enzymes; other enzymes; enzyme inhibitors; structural/signalling proteins; conserved uncharacterised proteins; and novel uncharacterised proteins [21].

We used the reference genomes of five echinoderms, including the Great Barrier Reef genome of COTS, which were downloaded from NCBI (*Strongylocentrotus purpuratus*) [64] and *Apostichopus japonicus* [65], Echinobase (*Asterias rubens* and *Patiria miniata)* [66], and https://marinegenomics.oist.jp/cots/viewer/info?project_id=46 (COTS) [21] to identify lineage-specific genes. Orthofinder was used with default settings [67] to identify orthologous groups, gene duplication events and putative species-specific coding sequences. COTS coding sequences that did not appear to share orthologues with other echinoderms or that had duplicates (paralogues) were considered to be lineage-specific. A Venn diagram of the number of shared and lineage-specific orthologous groups of sequences was generated using https://bioinformatics.psb.ugent.be/webtools/Venn/. A species tree was generated using Orthofinder [67].

### Differential gene expression analyses

To determine potential read count errors associated with lowly expressed genes [68], we tested different expression threshold levels of an average of ≥ 0.25 and ≥ 1 read per gene per library for all replicates of a given tissue and opted for the expression threshold of ≥ 0.25 mean reads per tissue to increase the chance of detecting lowly expressed coding sequences (S3 Fig) [28]. Using Wald Test statistics in the DESeq2 package (v1.28.1) [69], coding sequences expression levels with an adjusted *p*-value < 0.05 between conditions (tissues, season and sex) were identified as being significantly differentially expressed. We conducted pairwise DESeq2 analyses between all tissues, and coding sequences that were consistently upregulated in any given tissue were identified as being tissue-specific. Similarly, for each tissue, the summer male and female groups were individually tested against the winter individuals using DESeq2. Coding sequences that were upregulated in both the male and female group compared to winter, were considered upregulated in summer, and vice versa.

To detect modules of co-expressed genes from the transcriptome data, Weighted Gene Correlation Network Analysis (WGCNA) was applied to normalised VST data [40]. Coding sequences expressed at low levels and with low expression variance across the libraries were filtered out. The remaining 9,159 coding sequences were used in the WGCNA analysis. A signed network was constructed in WGCNA with specific parameter settings of power = 7, TOMType = ‘signed Nowick’ and minModuleSize = 100 [40]. Networks significantly correlated with the summer sampling were visualised in Cytoscape (v. 3.9.1) [70] and were filtered based on edge weight.

### Enrichment analyses and data visualisation

Gene Ontology (GO) enrichment analyses of coding sequences were performed using Blast2GO as previously described [29], using the Fisher’s exact test function available on the “clusterProfiler” package (v4.2.2) [71]. GO terms with a False Discovery Rate (FDR) adjusted *p*-value of < 0.05 were considered enriched. GO enrichment analysis for the lineage-specific genes was performed against all the genes in the COTS GBR v1.1 genome [29], whereas GO enrichment analyses of tissue-specific or seasonally-upregulated genes were performed against the genes expressed in specific tissues. KOBAS-i was used to test the statistical enrichment of KEGG pathways in differentially expressed coding sequences, with a FDR-corrected *p*-value of < 0.05 [72]. Genes expressed in each tissue were used as reference for the enrichment analyses. GO and KEGG enrichment analyses were visualised using the “ggplot2” R package (v3.3.6) [73].

Principal component analyses (PCAs) were used to visualise differences in gene expression between tissues and seasons. PCAs were performed on variance-stabilising transformed (VST) counts obtained with DESeq2. Heatmaps were created for each tissue separately, to visualise expression patterns of genes of interest, using transcripts per million (TPM) normalised raw reads, in the R package “pheatmap” [74,75].

All analyses and visualisations were performed in RStudio version 4.0.2 [76].

## Acknowledgments

This research was support by a Linkage Project grant (LP170101049) from the Australian Research Council. We thank the Great Barrier Reef Foundation and Association of Marine Park Tourism Operators Limited (AMPTO) for their financial and logistical support; the crew and divers of the AMPTO vessels that assisted in the collection of COTS from Lynchs and Davies Reefs; and Nick Rhodes, Michael Thang, Gareth Price and Dominique Gorse from the Queensland Cyber Infrastructure Foundation for development of the COTS genome browser and computing assistance. DJC and CKW are supported by a Fellowship from the National Health and Medical Research Council, Australia (2009564) and by access to the facilities of the Australian Research Council Centre of Excellence for Innovations in Peptide and Protein Science (CE200100012) and an ARC Future Fellow (FFT220100583).

## Author Contributions

**Conceptualization:** Bernard M. Degnan, Sandie M. Degnan, Marie Morin, Mathias Jönsson.

**Data curation:** Marie Morin, Mathias Jönsson.

**Formal analysis:** Marie Morin, Mathias Jönsson.

**Funding acquisition:** Bernard M. Degnan, David J. Craik, Sandie M. Degnan, Conan K. Wang.

**Investigation:** Marie Morin, Mathias Jönsson, Bernard M. Degnan, Sandie M. Degnan.

**Methodology:** Marie Morin, Mathias Jönsson, Bernard M. Degnan, Sandie M. Degnan.

**Project administration:** Bernard M. Degnan, Sandie M. Degnan

**Resources:** Bernard M. Degnan, Sandie M. Degnan, Marie Morin, Mathias Jönsson.

**Supervision:** Bernard M. Degnan, Sandie M. Degnan

**Visualization:** Marie Morin, Mathias Jönsson, Bernard M. Degnan, Sandie M. Degnan.

**Writing – original draft:** Marie Morin, Mathias Jönsson, Bernard M. Degnan, Sandie M. Degnan.

**Writing – review & editing:** Bernard M. Degnan, Sandie M. Degnan, Marie Morin, Mathias Jönsson, David J. Craik, Conan K. Wang.

## Supporting information

**S1 Fig. Identification and expression of COTS-specific genes. (A)** Venn diagram showing the overlap of orthologous gene groups among five echinoderms, the sea urchin *Strongylocentrotus purpuratus* [64], sea cucumber *Apostichopus japonicus* [65], and starfish *Asterias rubens*, *Patiria miniata* [66] and *Acanthaster planci* cf. *solaris* (COTS) [21,29]. COTS has 506 unique gene groups. **(B)** Phylogenetic tree of the five echinoderms showing gene duplication events at each node and branch. The taxonomic ranks are listed on the right. **(C)** GO term enrichments of COTS-specific coding sequences. The x-axis shows the GO terms and the y-axis shows the number of coding sequences in each term. **(D, E)** Expression profiles of lineage-specific GPCRs and secreted proteins in wild COTS. The heatmaps show the scaled expression levels (z-score) based on TPM normalized reads for each tissue in seven females, six males and seven winter individuals. The “All” heatmap shows the average expression across all groups and tissues. The heatmaps are color-coded (right) by differential expression: non-differentially expressed (brown), summer-upregulated (yellow) and winter-upregulated (blue).

**S2 Fig. WGCNA module identification and correlation analysis. (A)** Clustering dendrogram of all samples. The colour bar at the bottom corresponds to COTS tissues: green, radial nerve; pink, tube feet; purple, sensory tentacles; brown, papulae; turquoise, skin; blue, spines; red, coelomocytes. **(B)** Scale-free topology model fit for different soft-thresholding powers (left). The signed R^2^ value measures how well the scale-free topology assumption fits the observed network connectivity distribution. Mean network connectivity for different soft-thresholding powers (right). The mean connectivity is the average number of connections (edges) per node (gene) in the network for a given power value. **(C)** Module-trait associations. Each column represents a trait (tissue/season/sex), and each row represents an identified module. Red and blue colour notes positive and negative correlation with the trait, respectively. Numbers in each cell represent the correlation coefficient, and the *p*-value (in brackets).

**S3 Fig. Comparison of expression threshold of seasonal transcriptomes. (A)** PCA of tissue transcriptomes with an average expression threshold ≥0.25. **(B)** PCA of tissue transcriptomes with an average expression threshold ≥1. 95% confidence ellipses shown. Pink, tube feet; purple, sensory tentacles; green, radial nerve; turquoise, skin; brown, papulae; blue, spines; red, coelomocytes.

**S1 Table. Mapping rates of CEL-Seq2 transcriptome reads to the COTS GBR v1.1 genome using Bowtie2.**

**S2 Table. Expression profiles of coding sequences (CDS) in different tissues of the crown-of-thorns starfish (COTS).** The table shows the presence (1) or absence (0) of reads for each CDS in seven tissues: coelomocytes (Co), skin (Sk), spines (Sp), papulae (Pa), radial nerve (RN), tube feet (TF), and sensory tentacles (ST). The table also provides information on the function, annotation, and classification of each CDS based on various databases and sources.

**S3 Table. Significantly upregulated coding sequences (CDS) in each tissue.**

**S3.1 Table.** Coelomocytes. **S3.2 Table.** Papulae. **S3.3 Table.** Radial nerve. **S3.4 Table.** Skin. **S3.5 Table.** Spines. **S3.6 Table.** Sensory tentacles. **S3.7 Table.** Tube feet. **S3.8 Table.** Significant gene ontology (GO) enrichments based on upregulated CDS in each tissue.

**S4 Table. Significantly upregulated coding sequences (CDS) in oral and aboral tissues.**

**S4.1 Table.** List of upregulated (1) and not upregulated (0) CDS in oral and aboral tissues. **S4.2 Table.** Significant gene ontology (GO) enrichments based on upregulated CDS in oral and aboral tissues.

**S5 Table. Differential expression analysis of coding sequences (CDS) in different tissues of the crown-of-thorns starfish (COTS) across seasons.**

**S5.1 Table.** DESeq2 was applied pairwise comparing summer and winter expression for each tissue. Upregulation (1) and no significant upregulation (0) was depicted across all tissues. **S5.2 Table.** Significant gene ontology (GO) enrichments based on upregulated CDS in summer and winter for all tissues. **S5.3 Table.** KEGG enrichments based on upregulated CDS in summer and winter for all tissues. **S5.4 Table.** Transcription factors significantly upregulated in the summer. **S5.5 Table.** Transcription factors significantly upregulated in the winter.

**S6 Table. Lineage-specific coding sequences that are differentially expressed in COTS somatic tissues.**

**S6.1 Table.** Number of differentially expressed lineage-specific coding sequences in each tissue. **S6.2 Table.** All differentially expressed lineage-specific coding sequences, across both season and sex. Samples were compared pairwise, female (F) summer vs male (M) summer; F summer vs winter; and M vs winter, where 1 depicts upregulation. **S6.3 Table.** Significant GO enrichments based on differentially-expressed lineage-specific coding sequences.

**S7 Table. Co-expressed coding sequences in cyan, pink and turquoise co-expression modules.**

**S7.1 Table.** Co-expressed coding sequence in the cyan module **S7.2 Table.** Significant GO enrichments based on CDS in the cyan module. **S7.3 Table.** Co-expressed coding sequence in the pink module **S7.4 Table.** Significant GO enrichments based on CDS in the pink module. **S7.5 Table.** Co-expressed coding sequence in the turquoise module **S6.6 Table.** Significant GO enrichments based on CDS in the turquoise module.

## References

1. Haszprunar G, Vogler C, Wörheide G. Persistent gaps of knowledge for naming and distinguishing multiple species of crown-of-thorns-seastar in the *Acanthaster planci* Species Complex. Diversity. 2017;9: 22. doi:10.3390/d9020022

2. Haszprunar G, Spies M. An integrative approach to the taxonomy of the crown-of-thorns starfish species group (Asteroidea: *Acanthaster*): A review of names and comparison to recent molecular data. Zootaxa. 2014;3841: 271–14. doi:10.11646/zootaxa.3841.2.6

3. Lucas JS. Crown-of-thorns starfish. Current Biology. 2013;23: R945–R946. doi:10.1016/j.cub.2013.07.080

4. Pratchett M, Caballes C, Wilmes J, Matthews S, Mellin C, Sweatman H, et al. Thirty years of research on crown-of-thorns starfish (1986–2016): scientific advances and emerging opportunities. Diversity. 2017;9: 41–49. doi:10.3390/d9040041

5. Yasuda N, Inoue J, Hall MR, Nair MR, Adjeroud M, Fortes MD, et al. Two hidden mtDNA-clades of crown-of-thorns starfish in the Pacific Ocean. Front Mar Sci. 2022;9: 831240. doi:10.3389/fmars.2022.831240

6. Allen JD, Richardson EL, Deaker D, Agüera A, Byrne M. Larval cloning in the crown-of-thorns sea star, a keystone coral predator. Marine Ecology Progress Series. 2019;609: 271–276. doi:10.3354/meps12843

7. Babcock RC, Dambacher JM, Morello EB, Plagányi ÉE, Hayes KR, Sweatman HPA, et al. Assessing different causes of crown-of-thorns starfish outbreaks and appropriate responses for management on the Great Barrier Reef. Plos One. 2016;11: e0169048. doi:10.1371/journal.pone.0169048

8. Babcock RC, Milton DA, Pratchett MS. Relationships between size and reproductive output in the crown-of-thorns starfish. Mar Biol. 2016;163: 234. doi:10.1007/s00227-016-3009-5

9. Babcock, R. C., Plagányi, É. E., Condie, S. A., Westcott, D. A., Fletcher, C. S., Bonin, M. C., & Cameron, D. (2020). Suppressing the next crown-of-thorns outbreak on the Great Barrier Reef. Coral Reefs, 39(5), 1233–1244. doi:10.1007/s00338-020-01978-8

10. Caballes CF, Byrne M, Messmer V, Pratchett MS. Temporal variability in gametogenesis and spawning patterns of crown-of-thorns starfish within the outbreak initiation zone in the northern Great Barrier Reef. Mar Biol. 2021;168: 13. doi:10.1007/s00227-020-03818-3

11. Deaker DJ, Byrne M. Crown of thorns starfish life-history traits contribute to outbreaks, a continuing concern for coral reefs. Emerg Top Life Sci. 2022;6: 67–79. doi:10.1042/etls20210239

12. Deaker DJ, Mos B, Lin H-A, Lawson C, Budden C, Dworjanyn SA, et al. Diet flexibility and growth of the early herbivorous juvenile crown-of-thorns sea star, implications for its boom-bust population dynamics. Russell BD, editor. PLoS ONE. 2020;15: e0236142–17. doi:10.1371/journal.pone.0236142

13. Pratchett M, Dworjanyn S, Mos B, Caballes C, Thompson C, Blowes S. Larval survivorship and settlement of crown-of-thorns starfish (*Acanthaster* cf. *solaris*) at varying algal cell densities. Diversity. 2017;9: 2–11. doi:10.3390/d9010002

14. Uthicke S, Schaffelke B, Byrne M. A boom–bust phylum? Ecological and evolutionary consequences of density variations in echinoderms. Ecol Monogr. 2009;79: 3–24. doi:10.1890/07-2136.1

15. Vercelloni J, Caley MJ, Mengersen K. Crown-of-thorns starfish undermine the resilience of coral populations on the Great Barrier Reef. Global Ecol Biogeogr. 2017;26: 846–853. doi:10.1111/geb.12590

16. Wolfe K, Graba-Landry A, Dworjanyn SA, Byrne M. Larval starvation to satiation: influence of nutrient regime on the success of *Acanthaster planci*. Ferse SCA, editor. PLoS ONE. 2015;10: e0122010–17. doi:10.1371/journal.pone.01220

17. Babcock R, Mundy C. Reproductive biology, spawning and field fertilization rates of *Acanthaster planci*. Mar Freshw Res. 1992;43: 525–533. doi:10.1071/MF9920525

18. Beach DH, Hanscomb NJ, Ormond RFG. Spawning pheromone in crown-of-thorns starfish. Nature. 2004;254: 135–136. doi:10.1038/254135a0

19. Caballes CF, Pratchett MS. Environmental and biological cues for spawning in the crown-of-thorns starfish. PLoS ONE. 2017;12: e0173964. doi:10.1371/journal.pone.0173964

20. Mercier A, Hamel JF. Endogenous and exogenous control of gametogenesis and spawning in echinoderms. Adv Mar Biol. 2009;55: 1–302. doi:10.1016/s0065-2881(09)55001-8

21. Hall MR, Kocot KM, Baughman KW, Fernandez-Valverde SL, Gauthier MEA, Hatleberg WL, et al. The crown-of-thorns starfish genome as a guide for biocontrol of this coral reef pest. Nature. 2017;544: 231–234. doi:10.1038/nature22033

22. Anderson WD, Soh JY, Innis SE, Dimanche A, Ma L, Langefeld CD, et al. Sex differences in human adipose tissue gene expression and genetic regulation involve adipogenesis. Genome Res. 2020;30: 1379–1392. doi:10.1101/gr.264614.120

23. Campbell-Staton SC, Velotta JP, Winchell KM. Selection on adaptive and maladaptive gene expression plasticity during thermal adaptation to urban heat islands. Nat Commun. 2021;12: 6195. doi:10.1038/s41467-021-26334-4

24. Jeffries KM, Teffer A, Michaleski S, Bernier NJ, Heath DD, Miller KM. The use of non-lethal sampling for transcriptomics to assess the physiological status of wild fishes. Comp Biochem Physiol Part B: Biochem Mol Biol. 2021;256: 110629. doi:10.1016/j.cbpb.2021.110629

25. Nanda S, Jacques MA, Wang W, Myers CL, Yilmaz LS, Walhout AJ. Systems-level transcriptional regulation of *Caenorhabditis elegans* metabolism. Mol Syst Biol. 2023;19:e11443., doi:10.15252/msb.202211443

26. Wucher V, Sodaei R, Amador R, Irimia M, Guigó R. Day-night and seasonal variation of human gene expression across tissues. PLOS Biol. 2023;21: e3001986. doi:10.1371/journal.pbio.3001986

27. Zhang W, Liu Z, Tang S, Li D, Jiang Q, Zhang T. Transcriptional response provides insights into the effect of chronic polystyrene nanoplastic exposure on *Daphnia pulex*. Chemosphere. 2020;238: 124563. doi:10.1016/j.chemosphere.2019.124563

28. Jönsson M, Morin M, Wang CK, Craik DJ, Degnan SM, Degnan BM. Sex-specific expression of pheromones and other signals in gravid starfish. BMC Biol. 2022;20: 288. doi:10.1186/s12915-022-01491-0

29. Morin M, Jönsson M, Wang CK, Craik DJ, Degnan SM, Degnan BM. Captivity induces a sweeping and sustained genomic response in a starfish. Mol Ecol. 2023;32: 3541–3556. doi:10.1111/mec.16947

30. Burke RD. Podial sensory receptors and the induction of metamorphosis in echinoids. J Exp Mar Biol Ecol. 1980;47: 223–234. doi:10.1016/0022-0981(80)90040-4

31. Elia L, Selvakumaraswamy P, Byrne M. Nervous system development in feeding and nonfeeding asteroid larvae and the early juvenile. Biol Bull. 2009;216: 322–334. doi:10.1086/BBLv216n3p322

32. Roberts RE, Motti CA, Baughman KW, Satoh N, Hall MR, Cummins SF. Identification of putative olfactory G-protein coupled receptors in crown-of-thorns starfish, *Acanthaster planci*. BMC Genomics. 2017;18: 1–15. doi:10.1186/s12864-017-3793-4

33. Roberts RE, Powell D, Wang T, Hall MH, Motti CA, Cummins SF. Putative chemosensory receptors are differentially expressed in the sensory organs of male and female crown-of-thorns starfish, *Acanthaster planci*. BMC Genomics. 2018;19: 1–13. doi:10.1186/s12864-018-5246-0

34. Sloan N. The arm curling and terminal tube-foot responses of the asteroid *Crossaster papposus* (L.). J Nat Hist. 1980;14:469–482. doi:10.1080/00222938000770411

35. Cobb J. An ultrastructural study of the dermal papulae of the starfish, *Asterias rubens*, with special reference to innervation of the muscles. Cell Tissue Res. 1978;187: 515–523. doi:10.1007/BF00229616

36. Brusca RC, Moore W, Shuster SM. Invertebrates. 3^rd^ Edition. 2016; Sinauer Associates Inc.

37. Guschina IA, Harwood JL. Mechanisms of temperature adaptation in poikilotherms. FEBS Lett. 2006;580: 5477–5483. doi:10.1016/j.febslet.2006.06.066

38. Rissanen E, Tranberg HK, Sollid J, Nilsson GE, Nikinmaa M. Temperature regulates hypoxia-inducible factor-1 (HIF-1) in a poikilothermic vertebrate, crucian carp (*Carassius carassius*). J Exp Biol. 2006;209: 994–1003. doi:10.1242/jeb.02103

39. Guerra V, Haynes G, Byrne M, Yasuda N, Adachi S, Nakamura M, et al. Nonspecific expression of fertilization genes in the crown-of-thorns *Acanthaster* cf. *solaris*: Unexpected evidence of hermaphroditism in a coral reef predator. Molecular Ecology. 2019;29: 363–379. doi:10.1111/mec.15332

40. Zhang B, Horvath S. (2005). A general framework for weighted gene co-expression network analysis. Statistical Applications in Genetics and Molecular Biology. 2005:4: 17. doi:10.2202/1544-6115.1128

41. Ariyoshi M, Schwabe JWR. A conserved structural motif reveals the essential transcriptional repression function of Spen proteins and their role in developmental signaling. Genes Dev. 2003;17: 1909–1920. doi:10.1101/gad.266203

42. Watson PJ, Fairall L, Schwabe JWR. Nuclear hormone receptor co-repressors: Structure and function. Mol Cell Endocrinol. 2012;348: 440–449. doi:10.1016/j.mce.2011.08.03

43. Levin ER, Hammes SR. Nuclear receptors outside the nucleus: extranuclear signalling by steroid receptors. Nat Rev Mol Cell Biol. 2016;17: 783–797. doi:10.1038/nrm.2016.122

44. Vihervaara A, Duarte FM, Lis JT. Molecular mechanisms driving transcriptional stress responses. Nat Rev Genet. 2018;19: 385–397. doi:10.1038/s41576-018-0001-6

45. Hotamisligil GS, Davis RJ. Cell signaling and stress responses. CSH Perspect Biol. 2016;8: a006072. doi:10.1101/cshperspect.a006072

46. Baird NA, Turnbull DW, Johnson EA. Induction of the heat shock pathway during hypoxia requires regulation of heat shock factor by hypoxia-inducible factor-1*. J Biol Chem. 2006;281: 38675–38681. doi:10.1074/jbc.m608013200

47. Klumpen E, Hoffschröer N, Zeis B, Gigengack U, Dohmen E, Paul RJ. Reactive oxygen species (ROS) and the heat stress response of *Daphnia pulex*: ROS-mediated activation of hypoxia-inducible factor 1 (HIF-1) and heat shock factor 1 (HSF-1) and the clustered expression of stress genes. Biol Cell. 2017;109: 39–64. doi:10.1111/boc.201600017

48. Roberts RJ, Agius C, Saliba C, Bossier P, Sung YY. Heat shock proteins (chaperones) in fish and shellfish and their potential role in relation to fish health: a review. J Fish Dis. 2010;33: 789–801. doi:10.1111/j.1365-2761.2010.01183.x

49. Otto JJ, Kane RE, Bryan J. Formation of filopodia in coelomocytes: localization of fascin, a 58,000 dalton actin cross-linking protein. Cell 1979;17: 285–293. doi:10.1016/0092-8674(79)90154-5

50. Edds KT. (1993). Cell biology of echinoid coelomocytes: I. Diversity and characterization of cell types. J Invert Path 1993; 61: 173–178. doi:10.1006/jipa.1993.1031

51. Wahltinez SJ, Byrne M, Stacy NI. Coelomic fluid of asteroid echinoderms: Current knowledge and future perspectives on its utility for disease and mortality investigations. Vet Path. 2023;60(5):547–559. doi:10.1177/03009858231176563

52. Watts HE. Seasonal regulation of behaviour: what role do hormone receptors play? Proc R Soc B: Biol Sci. 2020;287: 20200722. doi:10.1098/rspb.2020.0722

53. Voogt PA, Besten PJ den, Jansen M. Steroid metabolism in relation to the reproductive cycle in *Asterias rubens* L. Comp Biochem Physiol Part B: Comp Biochem. 1991;99: 77–82. doi:10.1016/0305-0491(91)90010-b

54. Thongbuakaew T, Suwansa-ard S, Chaiyamoon A, Cummins SF, Sobhon P. Sex steroids and steroidogenesis-related genes in the sea cucumber, *Holothuria scabra* and their potential role in gonad maturation. Sci Rep. 2021;11: 2194. doi:10.1038/s41598-021-81917-x

55. Hafez MSMAE, Okbah MAEA, Ibrahim HAH, Hussein AAER, Moneim NAAE, Ata A. First report of steroid derivatives isolated from starfish *Acanthaster planci* with anti-bacterial, anti-cancer and anti-diabetic activities. Nat Prod Res. 2022;36: 5545–5552. doi:10.1080/14786419.2021.2021200

56. Nelson HR, Altieri AH. Oxygen: the universal currency on coral reefs. Coral Reefs. 2019;38: 177–198. doi:10.1007/s00338-019-01765-0

57. Hashimshony T, Senderovich N, Avital G, Klochendler A, Leeuw Y de, Anavy L, et al. CEL-Seq2: sensitive highly-multiplexed single-cell RNA-Seq. Genome Biol. 2016;17: 1–7. doi:10.1186/s13059-016-0938-8

58. Andrews S. FastQC: a quality control tool for high throughput sequence data. Babraham Bioinformatics, Babraham Institute, Cambridge, United Kingdom; 2010.

59. Anders S, Pyl PT, Huber W. HTSeq—a Python framework to work with high-throughput sequencing data. Bioinform. 2014; 31: 166–169. doi:10.1093/bioinformatics/btu638

60. Sogabe S, Hatleberg WL, Kocot KM, Say TE, Stoupin D, Roper KE, et al. Pluripotency and the origin of animal multicellularity. Nature. 2019;570: 1–20. doi:10.1038/s41586-019-1290-4

61. Almagro Armenteros JJ, Tsirigos KD, Sønderby CK et al. SignalP 5.0 improves signal peptide predictions using deep neural networks. Nat Biotechnol 2019;37: 420–423. doi:10.1038/s41587-019-0036-z

62. Krogh A, Larsson B, Von Heijne G, Sonnhammer EL. Predicting transmembrane protein topology with a hidden Markov model: application to complete genomes. J Mol Biol. 2001;305: 567–580. doi:10.1006/jmbi.2000.4315

63. Voss, M., Schröder, B., & Fluhrer, R. (2013). Mechanism, specificity, and physiology of signal peptide peptidase (SPP) and SPP-like proteases. Biochim Biophys Acta -Biomembr. 2013;1828: 2828–2839. doi:10.1016/j.bbamem.2013.03.033

64. Sodergren E, Weinstock GM, Davidson EH, Cameron RA, Gibbs RA, et al. The genome of the sea urchin *Strongylocentrotus purpuratus*. Science. 2006;314: 941–952. doi:10.1126/science.1133609

65. Jo J, Oh J, Lee H-G, Hong H-H, Lee S-G, Cheon S, et al. Draft genome of the sea cucumber *Apostichopus japonicus* and genetic polymorphism among color variants. Gigascience. 2017;6:giw006. doi:10.1093/gigascience/giw006

66. Arshinoff BI, Cary GA, Karimi K, Foley S, Agalakov S, Delgado F, et al. Echinobase: leveraging an extant model organism database to build a knowledgebase supporting research on the genomics and biology of echinoderms. Nucleic Acids Res. 2021;50: D970–D979. doi:10.1093/nar/gkab1005

67. Emms DM, Kelly S. OrthoFinder: phylogenetic orthology inference for comparative genomics. Genome Biol. 2019;20: 238. doi:10.1186/s13059-019-1832-y

68. Y Sha, JH Phan, Wang MD. Effect of low-expression gene filtering on detection of differentially expressed genes in RNA-seq data. 37th Ann Intl Conf IEEE Eng Med Biol Soc. (EMBC), Milan, Italy, 2015; 6461–6464, doi:10.1109/EMBC.2015.7319872.

69. Love MI, Huber W, Anders S. Moderated estimation of fold change and dispersion for RNA-seq data with DESeq2. Genome Biol. 2014; 15: 1–21. doi:10.1186/s13059-014-0550-8

70. Su G, Morris JH, Demchak B, Bader GD. (2014). Biological network exploration with Cytoscape 3. Curr Protocol Bioinform. 2014;47: 8–13. doi:10.1002/0471250953.bi0813s47

71. Wu T, Hu E, Xu S, Chen M, Guo P, Dai Z, et al. clusterProfiler 4.0: A universal enrichment tool for interpreting omics data. Innovation, 2021; 2: 100141. doi:10.1016/j.xinn.2021.100141

72. Bu D, Luo H, Huo P, Wang Z, Zhang S, He Z, et al. KOBAS-i: intelligent prioritization and exploratory visualization of biological functions for gene enrichment analysis. Nucl Acids Res. 2021;49: W317–W325. doi:10.1093/nar/gkab447

73. Wickham H. ggplot2: elegant graphics for data analysis Springer-Verlag New York; 2009. Preprint at. 2016.

74. Kolde R. Package ‘pheatmap’. 2015; R package

75. Wagner GP, Kin K, Lynch VJ. Measurement of mRNA abundance using RNA-seq data: RPKM measure is inconsistent among samples. Theory Biosci. 2012;131: 281–285. doi:10.1007/s12064-012-0162-3

76. RStudio Team. RStudio: integrated development for R. 2020; RStudio, PBC, Boston, MA.

