## Supplementary figures and images for "Seasonal tissue-specific gene expression reveals reproductive and stress-related transcriptional systems in wild crown-of-thorns starfish"

### S1 Fig.pdf

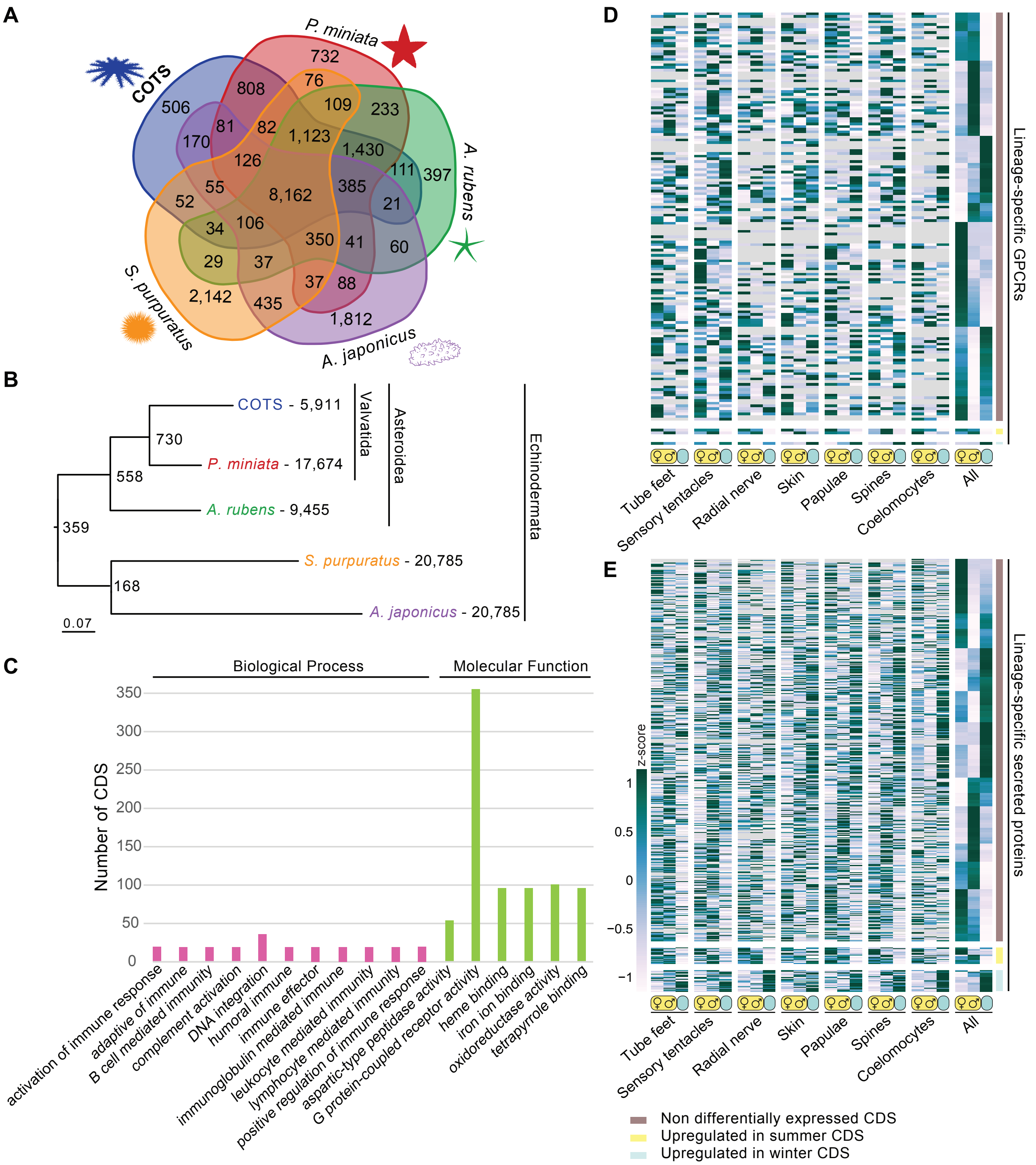

### S2 Fig.pdf

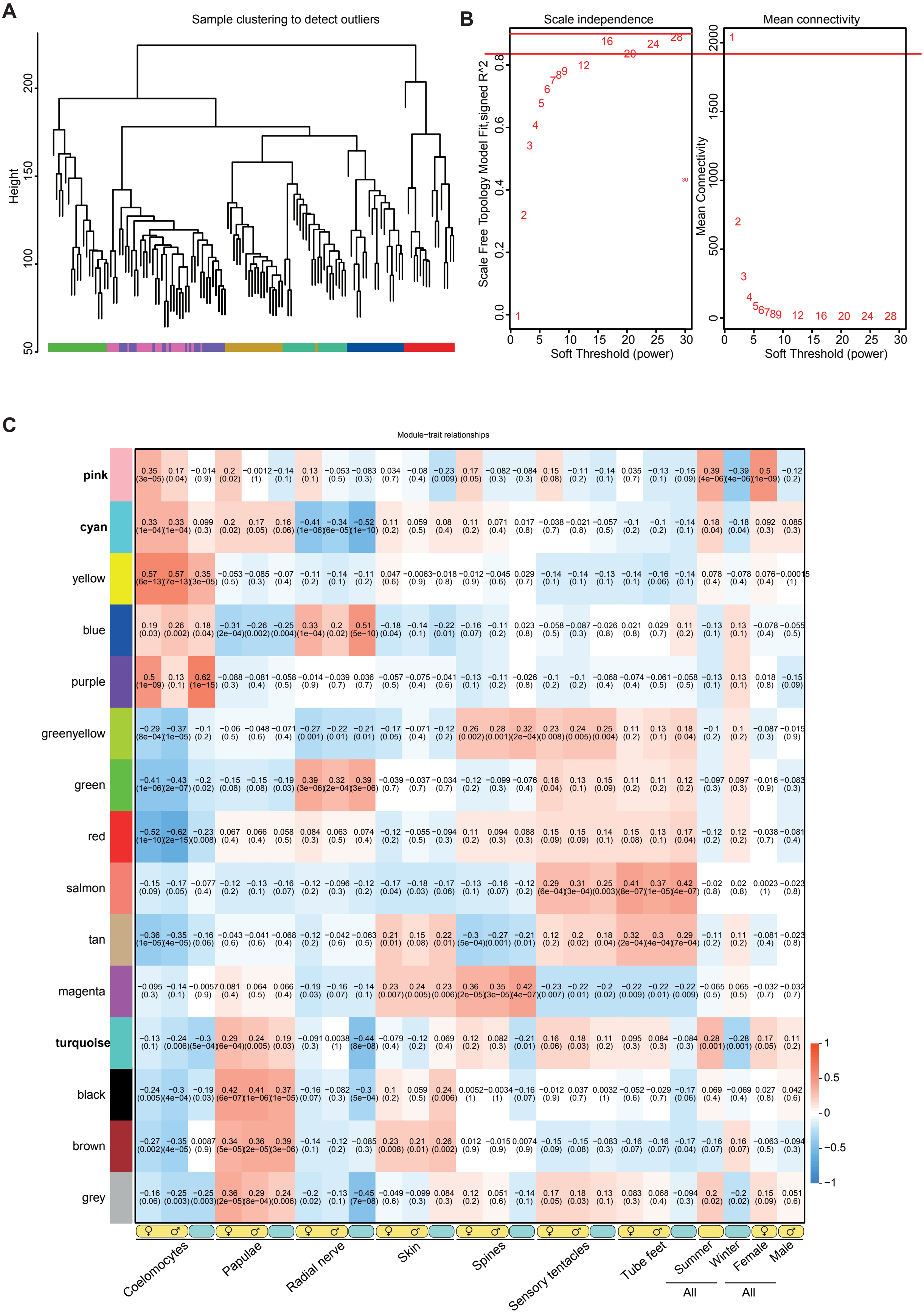

### S3 Fig.pdf

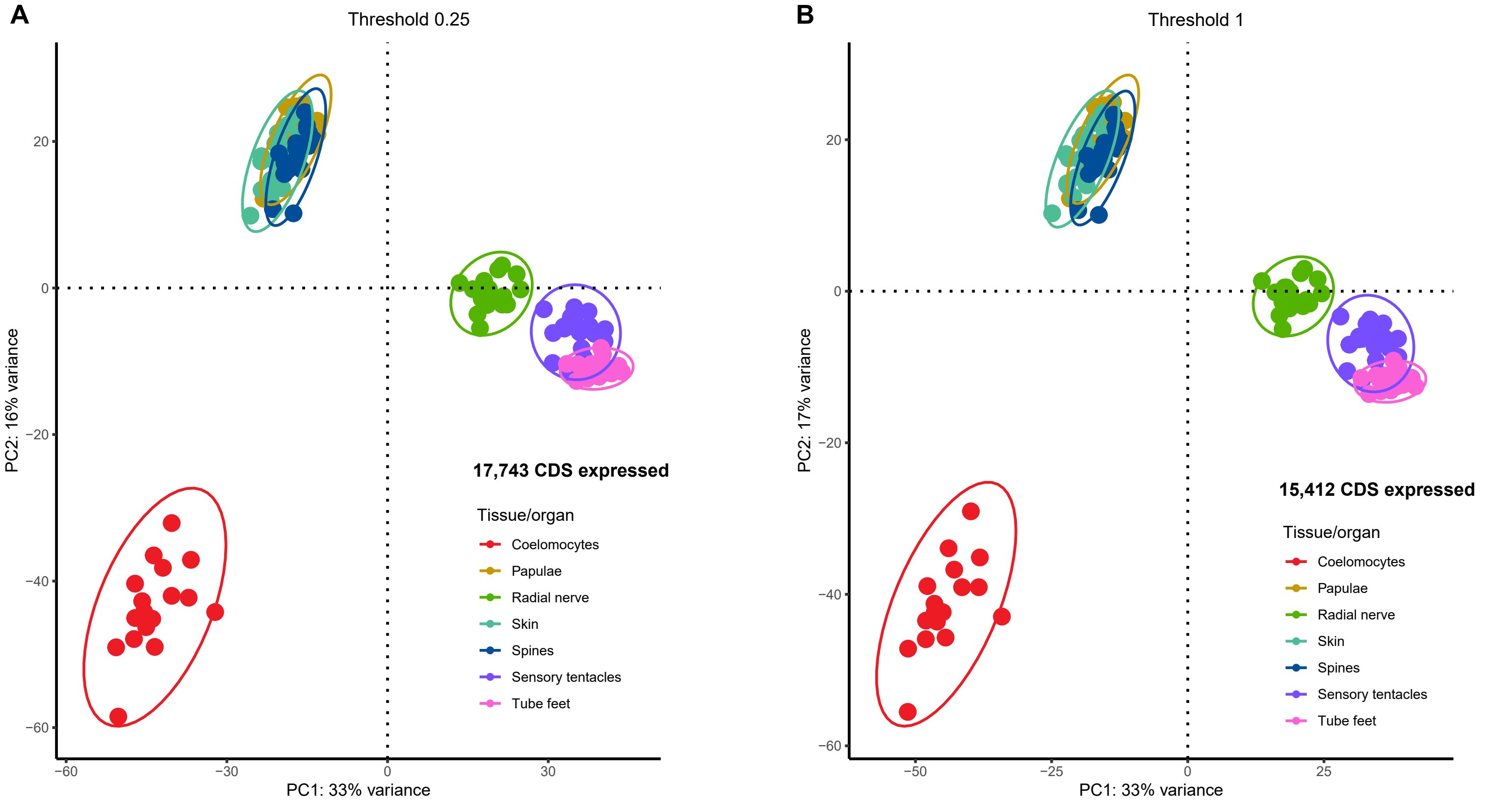
